# Host mucin is subverted by *Pseudomonas aeruginosa* during infection to provide free glycans required for successful colonization

**DOI:** 10.1101/675538

**Authors:** Casandra L Hoffman, Alejandro Aballay

## Abstract

The mucosal barrier, found lining epithelial cells, serves multiple functions in a range of animals. The major structural components of mucus are mucins, which are heavily glycosylated proteins that are either membrane bound or secreted by the epithelial cells. Mucins are key components of the innate immune system, as they are involved in the clearance of pathogens from the airways and intestines, and their expression is typically upregulated upon epithelial cell exposure to a variety of pathogens. In this study, we identified the mucin MUL-1 as an innate immune factor that appears to be utilized by *P. aeruginosa* to colonize hosts. We found that while the expression of several mucins, including MUL-1, increased upon *P. aeruginosa* infection of the nematode *Caenorhabditis elegans*, silencing of or deletion of *mul-1* resulted in enhanced survival and reduced bacterial accumulation. *P. aeruginosa* required host sialidase CTSA-1.1 to use mucin-derived glycans to colonize the host, while sialidase-encoding bacteria required host MUL-1 but not CTSA-1.1 to cause a lethal infection. This role of mucins and free glycans in host-pathogen interaction appears to be conserved from *C. elegans* to humans, as *P. aeruginosa* binding to human lung epithelial cells was also enhanced in the presence of free glycans, and free glycans reversed the binding defect of *P. aeruginosa* to human lung cells lacking the mucin MUC1.

**Author Summary:** The gastrointestinal, respiratory, reproductive, and urinary tracts, are large surfaces exposed to the exterior environment and thus, these mucosal epithelial tissues serve as primary routes of infection. One of the first lines of defense present at these barriers is mucus, which is a highly viscous material formed by mucin glycoproteins. Mucins serve various functions, but importantly they aid in the clearance of pathogens and debris from epithelial barriers and serve as innate immune effectors. In this study, we describe the ability of *Pseudomonas aeruginosa* to utilize mucin-derived glycans to colonize the intestine and ultimately cause death in *Caenorhabditis elegans*. We also show conserved mechanisms of *P. aeruginosa* virulence traits, by demonstrating that free glycans alter the ability of the bacteria to bind to human lung alveolar epithelial cells. Over the course of host-pathogen evolution, pathogens seem to have evolved to use mucins for their own advantage, and thus one of the biggest questions is which party benefits from pathogen-mucin binding. By gaining a better understanding of pathogen-mucin interactions, we can better protect against pathogen infection.

## Introduction

The mucus layer found lining mucosal epithelial cells is a crucial first barrier against microbial infections. The main structural components of the protective barrier of mucus are mucins, which are high molecular weight proteins, heavily glycosylated at serine and threonine residues (1–4). Mucins are secreted in large quantities by mucosal epithelial cells and both membrane-tethered and secreted mucins are found on the apical surface of all mucosal epithelium. Mucins aid in the formation of highly viscous, aqueous solutions that protect epithelial cells from physical damage, dehydration, and infection (1,2,5).

Epithelial barriers have evolved multiple mechanisms to respond to environmental cues, including those associated with pathogen exposure. At basal levels and in response to microbes, epithelial cells secrete defensive compounds, including mucins, defensins, lysozymes, and other antimicrobial compounds. Together, these compounds can form a physical barrier, have direct antimicrobial activity, and aid in pathogen clearance. Mucins are able to perform all of these activities. In addition to forming an impervious gel, mucus traps microbes, aids in the clearance of microbes, forms a physical barrier, and provides a matrix for a rich array of antimicrobial molecules (6,7).

Despite the various defensive properties of mucins, some bacterial pathogens are able to colonize mucosal epithelial barriers (6,7). Microbes must first penetrate the mucosal barrier to either attach to epithelial cells or release toxins that disrupt the epithelial barrier (7). One of the low-affinity binding mechanisms used by microbes is mediated by hydrophobic interactions with lectins and glycosylated receptors. Bacterial adhesins can bind to oligosaccharides present on mucins to mediate adhesion (8,9). One of the largest questions that remains to be answered is whether the bacterial-mucin binding event favors the bacteria or the host. It would appear that increased mucin secretion during infection would categorize mucins as a component of host defenses, but the coevolution of pathogens and hosts may allow for a range of roles for mucins for both parties. Some components of mucins may facilitate bacterial colonization of the host, including oligosaccharides that act as adhesion sites for bacteria and individual monosaccharides from mucins that can be accessed as energy sources by bacteria with mucolytic activity (10). Finally, mucus components can influence the virulence characteristics of pathogenic bacteria, such as virulence factor expression, adhesion, motility, proliferation, and/or growth (11–15).

Alterations in mucin expression or glycosylation patterns have been linked to various pathologies and diseases, including cystic fibrosis, chronic obstructive pulmonary disease, cancers, and inflammatory bowel disease (1). These diseases, that are linked to changes in mucin expression, glycosylation patterns, and alterations in mucus levels, are associated with higher rates of infection with opportunistic pathogens (2,16,17). Increases in mucus levels often result in a more highly viscous material that is not readily cleared from the mucosal barrier. Decreases in mucus levels remove the protective barrier, which allows pathogens to have direct access to epithelial cells. Many of the diseases linked to alterations in mucus levels have no known cure, and the primary mode of treatment involves controlling mucin expression, which is a key method used to prevent bacterial infections associated with these diseases (16,18). A better understanding of the mechanisms by which pathogens utilize mucins will provide novel approaches to control infectious processes and prevent bacterial colonization.

The complexity of mammalian epithelial mucosal surfaces and the additional functions of the innate immune system can mask the roles of individual mucins at epithelial barriers. Using the model organism, *Caenorhabditis elegans*, we can tease apart the roles of individual mucins at the intestinal epithelial barrier. This model provides the advantage of a simple organism’s small genome size and the absences of adaptive immunity. *C. elegans* eats bacteria found in decomposing organic matter (19). Several pathogens are present in the environment in which *C. elegans* feeds, which can impair the growth of the nematode and induce stress responses, ultimately leading to nematode death (20). The ability to distinguish between pathogenic and non-pathogenic microbes is critical for the survival of the nematodes and thus, *C. elegans* has methods to recognize and counteract pathogens. One of these responses is the upregulation of mucins during infection (21,22).

Herein we explore the role of *mul-1* during pathogen infection and find that pathogens use the immune factor to gain a competitive advantage in the host. We identified *mul-1* as a mucin, that when inhibited, enhances resistance to *P. aeruginosa* and *S. enterica* infection. This is despite the fact that *mul-1* gene expression is induced upon infection by these pathogens as part of the general immune response. We further explored the role of the mucin during infection and found that MUL-1 provides glycans that are required for attachment to *C. elegans* intestine and that a similar mechanism is conserved in mammalian epithelial cells. These results indicate that pathogens have evolved a mechanism to subvert conserved immune effectors to colonize different hosts.

## Results

### *C. elegans* lacking *mul-1* exhibit enhanced resistance to *P. aeruginosa* infection

Little is known about the roles of specific mucins in the *C. elegans* intestine during infection, but several enzymes and transporters that modify the glycan patterns of mucins have been identified to alter host-pathogen interactions. We set out to better understand the roles of specific intestine-expressed mucins during *P. aeruginosa* infection. The six identified mucins and mucin-editing enzymes (Table S1) were silenced by RNAi and the survival of the animals was monitored during *P. aeruginosa* PA14 infection. Upon analysis, RNAi for two genes, *mul-1* and *gpdh-1*, resulted in enhanced resistance to pathogen infection and RNAi for one mucin, *let-653*, resulted in enhanced susceptibility to pathogen infection (Table S1 and Figure S1).

Because mucins are upregulated upon infection and have been characterized as bona fide immune effectors, the enhanced resistance phenotype for two of the genes was unexpected. The most significant phenotype was observed for *mul-1* RNAi animals, which showed enhanced resistance to *P. aeruginosa* compared to RNAi control animals (Figure 1A). *mul-1* mRNA transcripts are found at high levels in the intestine (23) and it was previously shown that *mul-1* is upregulated upon infection with *P. aeruginosa* at 4 and 8 hours (21,22). Other upregulated genes from those datasets include innate immune factors, which would suggest that *mul-1* is an immune response factor turned on to combat pathogen infection.

**Figure 1.**
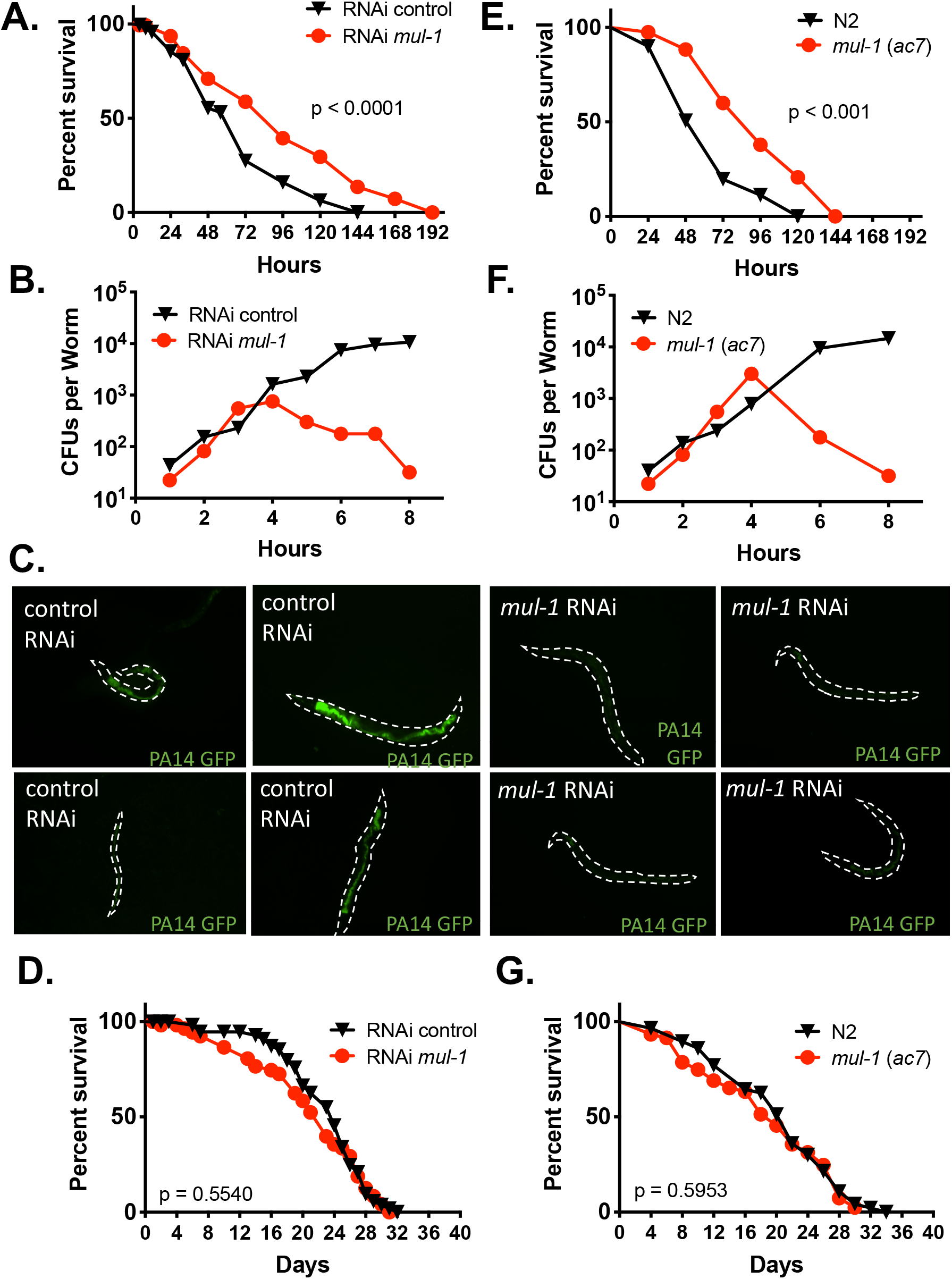
RNAi silencing and deletion of *mul-1* results in an enhanced resistance phenotype to *P. aeruginosa* PA14. **(A)** N2 wild-type animals were exposed to two generations of RNAi. Young adult animals were transferred to full lawns of *P. aeruginosa* PA14 and nematode survival was monitored daily. Animals were considered dead upon failure to respond to touch. Animals missing from the agar plate were censored on the day of loss. The KaplanMeier method was used to calculate the survival fractions, and statistical significance between survival curves was determined using the log-rank test. 3 biological replicates, 180 total animals per condition. P values compared to RNAi control: *mul-1* p < 0.0001, *let-653* p < 0.001, *gly-8* p = 0.5594, *cwp-4* p = 0.1336, *gpdh-1* p = 0.0079. **(B)** After RNAi, young adult nematodes were transferred to full lawns of *P. aeruginosa* PA14-GFP (kan^r^). At indicated time points, nematodes were transferred to fresh *E. coli* OP50 lawns to remove *P. aeruginosa*. Worms were ground and serial dilutions were plated on LB-kanamycin plates to calculate Colony Forming Units (CFUs) per worm (nematode). 3 biological replicates, 90 total animals per condition. **(C)** After RNAi, young adult nematodes were transferred to full lawns of *P. aeruginosa* PA14-GFP (kan^r^). At 24 hours, nematodes were transferred to fresh *E. coli* OP50 lawns to remove *P. aeruginosa*. Animals were anesthetized using an M9 salt solution containing 30 mM sodium azide and mounted onto 2% agar pads. The animals were visualized using a Leica M165 FC fluorescence stereomicroscope. Representative images of ~50 animals per condition from 2 biological replicates. **(D)** N2 wild-type animals were exposed to two generations of RNAi and L4 larval stage animals were transferred to fresh plates with the corresponding RNAi clone, heat-killed *E. coli* HT115(DE3). Survival was monitored daily. Worms were transferred to new plates as food was depleted. 3 biological replicates, 180 total animals per condition. **(E)** Young adult N2 wild-type and *mul-1 (ac7)* deletion animals were transferred to full lawns of *P. aeruginosa* PA14 and nematode survival was monitored daily. Animals were considered dead upon failure to respond to touch. Animals missing from the agar plate were censored on the day of loss. The KaplanMeier method was used to calculate the survival fractions, and statistical significance between survival curves was determined using the log-rank test. 3 biological replicates, 180 total animals per condition. **(F)** Young adult N2 wild-type and *mul-1(ac7)* deletion animals were transferred to full lawns of *P. aeruginosa* PA14-GFP (kan^r^). At indicated time points, nematodes were transferred to fresh *E. coli* OP50 lawns to remove *P. aeruginosa*. Worms were ground and serial dilutions were plated on LB-kanamycin plates to calculate Colony Forming Units (CFUs) per worm (nematode). 3 biological replicates, 90 total animals per condition. **(G)** L4 N2 wild-type and *mul-1 (ac7)* deletion animals transferred to fresh plates with heat-killed *E. coli* OP50. Survival was monitored daily. Worms were transferred to new plates as food was depleted. 3 biological replicates, 180 total animals per condition.

### The enhanced resistance of *mul-1* RNAi animals to *P. aeruginosa* is due to lack of bacterial colonization

Because *mul-1* RNAi results in delayed death upon *P. aeruginosa* infection, we aimed to better understand the mechanism by which *mul-1* contributes to enhanced resistance to the pathogen. We tested the ability of *P. aeruginosa* to colonize the nematode intestine. RNAi for *mul-1* resulted in reduced accumulation of bacteria, determined by obtaining *P. aeruginosa* Colony Forming Units (CFUs) from infected animals over the course of infection and measuring the fluorescence intensity in the intestines of animals infected with *P. aeruginosa* expressing GFP. Initially, RNAi *mul-1* animals accumulated *P. aeruginosa* at a similar rate as control animals, but there were significantly fewer CFUs recovered per nematode after 4 hours (Figure 1B) and little to no fluorescent signal was found in RNAi *mul-1* animals compared to control RNAi animals at 24 hours post infection (Figure 1C). To determine if the enhanced resistance to *P. aeruginosa* infection is a general phenotype in the *mul-1* RNAi animals, we exposed these animals to *Salmonella enterica* ser Typhimurium ST1344 and monitored survival. The enhanced resistance phenotype was also observed when the RNAi *mul-1* nematodes were exposed to full lawns of *S. enterica*. RNAi *mul-1* nematodes lived longer upon exposure to *S. enterica* than RNAi control nematodes (Supplementary Figure 1A) and accumulated fewer *S. enterica* bacteria (Supplementary Figure 1B and 1C).

Because of the increase in survival upon pathogen exposure observed in *mul-1* RNAi animals, we studied if differences in the lifespan of the *mul-1* RNAi animals may be responsible for the enhanced survival in the presence of *P. aeruginosa*. As shown in Figure 1D, the longevity of *mul-1* RNAi animals was indistinguishable from that of control animals, ruling out that the possibility that enhanced longevity was the cause of enhanced survival in the presence of pathogenic bacteria. To confirm the roles of *mul-1* during infection, an in-frame, full gene deletion strain was used. The *mul-1(ac7)* strain also showed enhanced resistance to *P. aeruginosa* (Figure 1E), significantly fewer CFUs per nematode (Figure 1F), and there was no difference in lifespan in comparison to wildtype N2 nematodes (Figure 1G).

Expression of *mul-1* has been reported in the intestine, hypodermis, and PVD and OLL neurons (23,24). To address whether intestinal *mul-1* is specifically contributing to infection, intestine-specific RNAi was performed using two intestine-specific RNAi strains, VP303 (25) and MGH171 (26). In both cases, we found that *mul-1* RNAi in the intestine resulted in enhanced resistance to *P. aeruginosa* (Figure 2A and 2B).

**Figure 2.**
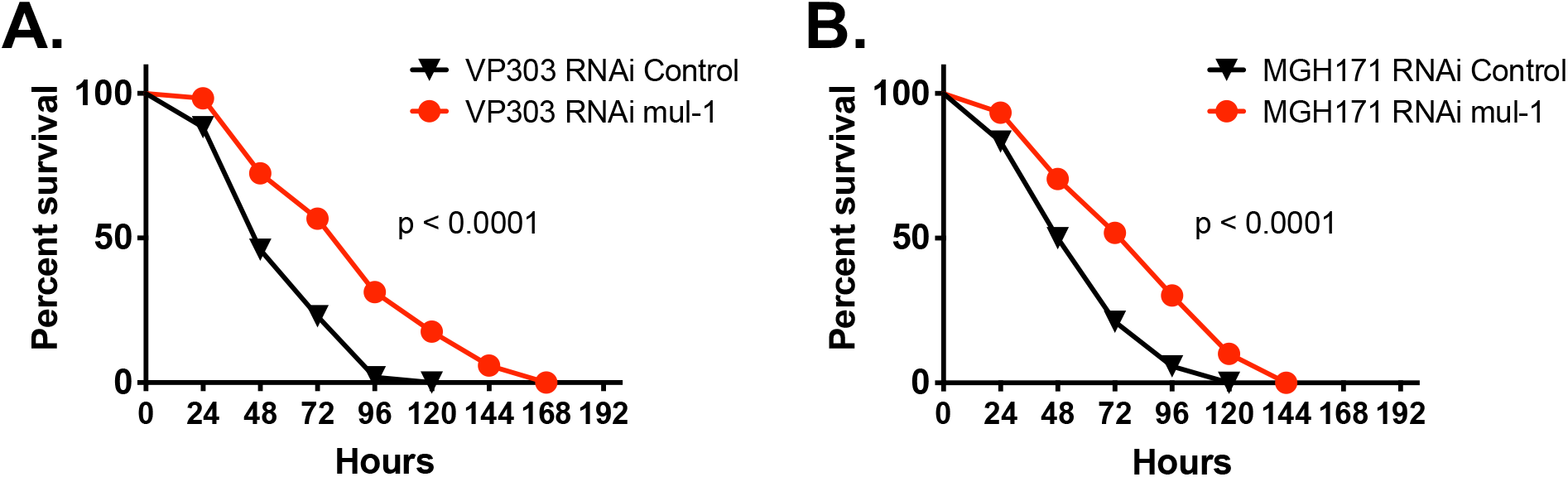
The enhanced resistance phenotype observed in *mul-1* silenced animals is intestine-specific. Intestine-specific RNAi animals, **(A)** VP303 and **(B)** MGH171, were exposed to two generations of RNAi. Young adult animals were transferred to full lawns of *P. aeruginosa* PA14 and nematode survival was monitored daily. Animals were considered dead upon failure to respond to touch. Animals missing from the agar plate were censored on the day of loss. The KaplanMeier method was used to calculate the survival fractions, and statistical significance between survival curves was determined using the log-rank test. 3 biological replicates, 180 animals per condition total.

One of the possible explanations for the *mul-1* RNAi animals’ enhanced resistance to *P. aeruginosa* could be due to the increased expression of other intestinal-expressed mucins that compensate for the loss of MUL-1. To rule out the possibility that other mucins are upregulated to compensate for the loss of *mul-1*, qRT-PCR was performed on RNAi control vector and RNAi *mul-1* nematodes. When the RNAi *mul-1* nematodes were grown on non-pathogenic *E. coli*, there were some statistically significant increases in expression levels of the intestine-expressed mucins and mucin-related enzymes compared to RNAi control nematodes. (Figure 3). Many of the significant increases found when the nematodes were grown on *E. coli* were no longer observed in *mul-1* RNAi nematodes grown on *P. aeruginosa* compared to control animals also exposed to *P. aeruginosa*. We don’t think these less than threefold increases could account for the enhanced resistance phenotype of the *mul-1* RNAi animals.

**Figure 3.**
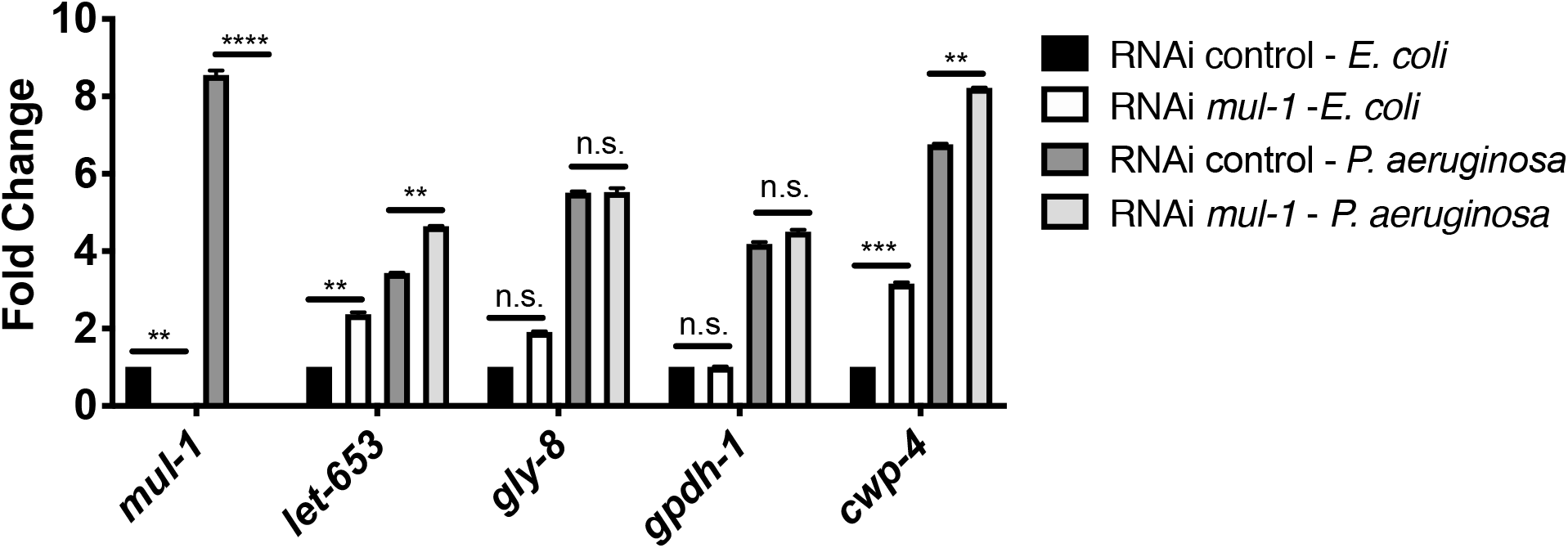
Silencing of *mul-1* by RNAi has little effect on the expression of other intestine-expressed mucins during infection. qRT-PCR analysis of the expression of *mul-1*, *let-653*, *gly-8*, *gpdh-1*, and *cwp-4* during growth on both *E. coli* OP50 and *P. aeruginosa* PA14 for 8 hours. N2 wild-type animals were exposed to two generations of RNAi. L4 larval stage animals were transferred to full lawns of *E. coli* OP50 or *P. aeruginosa* PA14 for 8 hours. RNA was isolated as described in methods section. Expression for indicated gene, under each RNAi condition, is compared to N2 wild-type RNAi control animals grown on *E. coli* OP50. 3 biological replicates, each with 3 technical replicates. ***p < 0.001 and **p < 0.01 via the t test. n.s., non-significant.

We have shown that *mul-1* RNAi animals are not colonized by *P. aeruginosa* during infection. Recent work shows that colonization by *P. aeruginosa* and other pathogens causes bloating of the intestine that results in the expression of innate immune effectors and the elicitation of a pathogen avoidance behavior (27). Because the *mul-1* mutant has limited bacterial accumulation in the intestine, we hypothesized that these animals would not bloat and thus would not avoid *P. aeruginosa*. *C. elegans* strains were exposed to partial lawns of *P. aeruginosa* PA14 and lawn occupancy was characterized at 24 hours. Consistent with the previous study, over 80% of the wild type N2 animals were outside of the *P. aeruginosa* PA14 lawn, demonstrating that wild type animals avoid *P. aeruginosa* (Figure 4A). In comparison, only 16.8% of the *mul-1 (ac7)* animals were outside of the bacterial lawn (83.2% lawn occupancy), suggesting that the mutant animals do not avoid *P. aeruginosa*. Because the *mul-1 (ac7)* animals do not avoid, it would be presumed that the animals have enhanced ingestion of *P. aeruginosa* and die more rapidly than wild type animals. Although the *mul-1 (ac7)* animals don’t avoid *P. aeruginosa*, they still demonstrate an enhanced resistance to the pathogen in comparison to wild type when survival was monitored on partial lawns of *P. aeruginosa* (Figure 4B).

**Figure 4.**
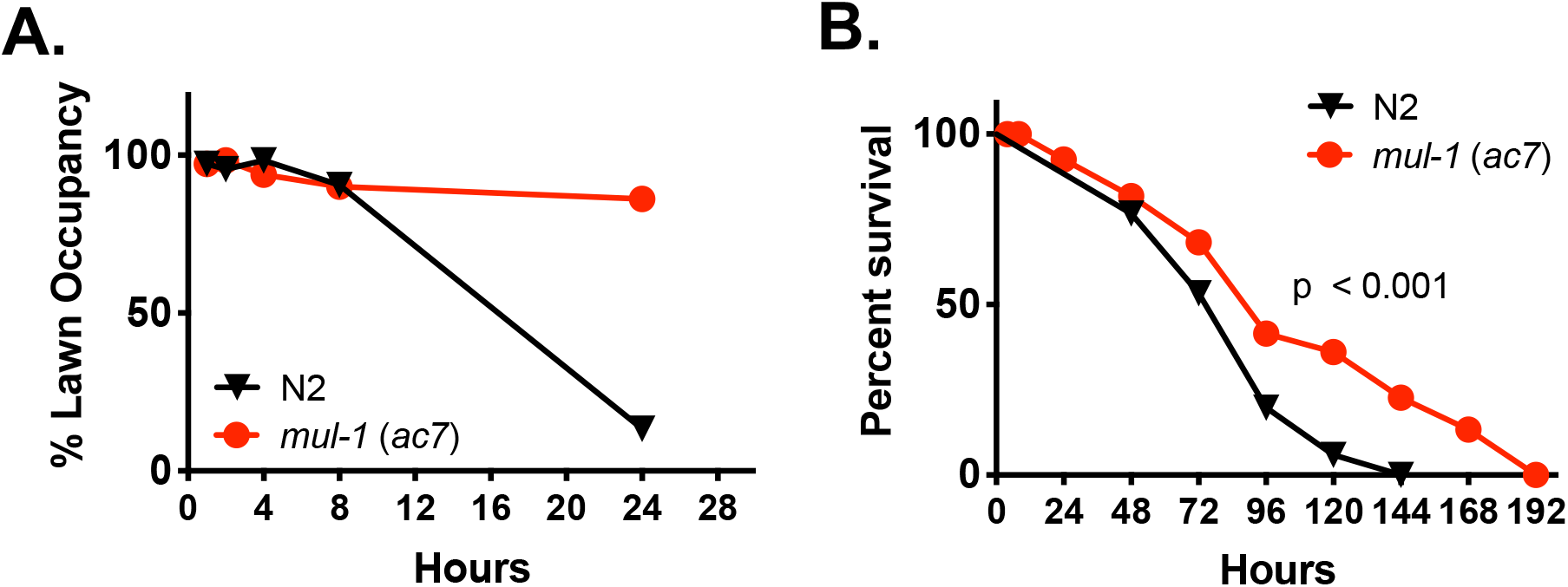
*mul-1* animals fail to avoid *P. aeruginosa* PA14 and failure to avoid has no effect on the enhanced resistance phenotype. **(A)** Young adult N2 wild-type and *mul-1 (ac7)* deletion animals were transferred to partial lawns of *P. aeruginosa* PA14. Percent (%) Lawn Occupancy was calculated at indicated time points as 100 × (the total number of animals inside the *P. aeruginosa* PA14 lawn / total number of animals outside the *P. aeruginosa* PA14 lawn). **(B)** Young adult N2 wild-type and *mul-1 (ac7)* deletion animals were transferred to partial lawns of *P. aeruginosa* PA14 and nematode survival was monitored daily. Animals were considered dead upon failure to respond to touch. Animals missing from the agar plate were censored on the day of loss. The KaplanMeier method was used to calculate the survival fractions, and statistical significance between survival curves was determined using the log-rank test. 3 biological replicates, 180 total animals per condition.

### Mucin-derived glycans are required for successful *P. aeruginosa* infection

Based on our data that shows *mul-1 (ac7)* and *mul-1* RNAi animals are not colonized by *P. aeruginosa*, we hypothesized that MUL-1 plays a beneficial role for pathogens in the host. This is contrary to the idea that MUL-1 is a *C. elegans* innate immune effector used by the nematode to fight and clear infections. To better understand the role of MUL-1 inside the *C. elegans* intestine, we probed whether *P. aeruginosa* is able to access glycans from MUL-1 in the *C. elegans* intestine.

In certain cases, pathogens have been reported to utilize glycans such as O-linked oligosaccharides from mucins (28–30), but in order to expose the host glycans, bacteria require a sialidase enzyme to cleave the sialic acid cap from host polysaccharides. *P. aeruginosa* does not encode a bacterial sialidase, thus we questioned if *C. elegans* expresses an enzyme that performs a similar function to process polysaccharides for *P. aeruginosa* use. *C. elegans* encodes CTSA-1.1, a homolog to human cathepsin A, which is predicted to have serine-type carboxypeptidase activity. This enzyme is predicted to have neuraminidase activity, which hydrolyzes terminal sialic acid residues on polysaccharide chains, most often exposing a galactose residue. The neuraminidase is expressed in the intestine, but does not change expression upon pathogen infection [21,31,32]. To determine if the neuraminidase contributes to the enhanced resistance to *P. aeruginosa* infection, RNAi for *ctsa-1.1* was performed. RNAi *ctsa-1.1* significantly enhanced resistance, but not to the same extent as *mul-1* RNAi (Figure 5A). Co-RNAi for both *mul-1* and *ctsa-1.1* was not additive, which suggests that these two gene products contribute to the same mechanism by which *P. aeruginosa* establishes an infection and kills *C. elegans*. There was no difference in PA14 accumulation in the RNAi *mul-1*, RNAi *ctsa-1.1*, or co-RNAi, but all three conditions resulted in significantly fewer CFUs per nematode compared to RNAi control (Figure 5B). To confirm that the enhanced resistance to *P. aeruginosa* phenotype was not due to enhanced longevity, we studied the *mul-1* RNAi, *ctsa-1.1* RNAi, and co-RNAi nematode lifespan. There were no differences in lifespan under all RNAi conditions (Figure 5C).

**Figure 5.**
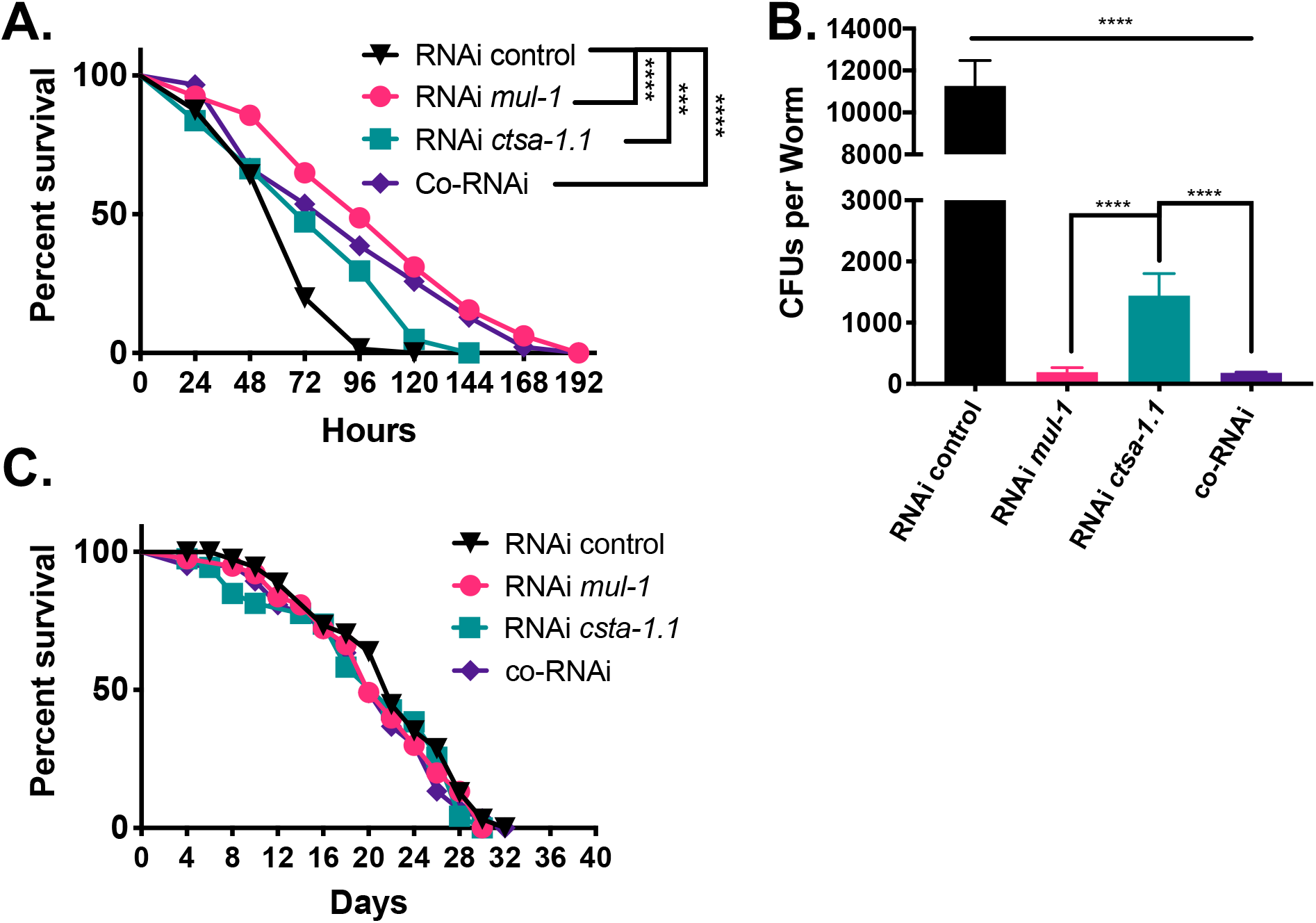
The ability of *P. aeruginosa* to access O-linked glycans provided by *mul-1* in the intestine alters the resistance phenotype of *C. elegans* nematodes. **(A)** N2 wild-type animals were exposed to two generations of RNAi targeting *mul-1*, *F41C3.5*, and an equal mixture of *mul-1* and *F41C3.5*. Young adult animals were transferred to full lawns of *P. aeruginosa* PA14 and nematode survival was monitored daily. Animals were considered dead upon failure to respond to touch. Animals missing from the agar plate were censored on the day of loss. The KaplanMeier method was used to calculate the survival fractions, and statistical significance between survival curves was determined using the log-rank test. 3 biological replicates, 180 total animals per condition. **(B)** After RNAi, young adult nematodes were transferred to full lawns of *P. aeruginosa* PA14-GFP (kan^r^). At 8 hours post *P. aeruginosa* PA14 exposure, nematodes were transferred to fresh *E. coli* OP50 lawns to remove *P. aeruginosa*. Worms were ground and serial dilutions were plated on LB-kanamycin plates to calculate Colony Forming Units (CFUs) per worm (nematode). 3 biological replicates, 90 total animals per condition. **(C)** N2 wild-type animals were exposed to two generations of RNAi and L4 larval stage animals were transferred to fresh plates with the corresponding RNAi clone, heat-killed *E. coli* HT115(DE3). Survival was monitored daily. Worms were transferred to new plates as food was depleted. 3 biological replicates, 180 total animals per condition.

Because the *mul-1* RNAi enhanced resistance phenotype was observed for both *P. aeruginosa* and *S. enterica*, we asked if both pathogens require the *C. elegans* neuraminidase, *ctsa-1.1*. Using the RNAi conditions above, nematodes were monitored for survival on *S. enterica*. As we have shown earlier, RNAi *mul-1* animals display an enhanced resistance phenotype when infected with *S. enterica*, but the RNAi *ctsa-1.1* animals are as susceptible to *S. enterica* as RNAi control, and accumulate as many *S. enterica* per nematode as control RNAi animals. (Supplementary Figure 2A-C). *S. enterica* encodes and expresses functional bacterial sialidases that allow the bacteria to access glycan molecules from mucins (31). Thus, the *C. elegans* enzyme may not be required by *S. enterica* during infection. The use of the *C. elegans* enzyme, CTSA-1.1 appears to be specific to *P. aeruginosa*, which lacks a sialidase.

To further elucidate the mechanism(s) by which mucins benefit pathogens during infection, we tested if individual purified free glycans had any effect on bacterial growth. We also wanted to determine if free glycans had any impact on *C. elegans* survival on *P. aeruginosa*. Based on previous studies that aimed to understand the impact of mucin glycosylation patterning on development and infectious disease, we identified several o-linked glycans that may be important for bacterial binding and or growth *in vivo* in the nematode. These included N-acetylgalactosamine, galactose, N-acetylglucosamine, and N-acetyl neuraminic acid, which is the predominant sialic acid found in mammalian mucin glycan chains (32–34). Commercially available free glycans were added to Luria Broth (LB) to assess their effect on PA14 growth. There were no significant changes to bacterial growth in LB (Supplemental Figure 3A). We also tested the effects of these free glycans in a minimal bacterial growth media, M9 media, because LB media is rich and provides ample carbon source(s) for the bacteria, and in LB media we may not observe any additional enhancement to growth. None of the free glycans had any effect on bacterial growth in M9 media over the course of 48 hours. In fact, we observed little to no growth of the bacteria in M9 media or M9 media supplemented with glycans (Supplemental Figure 3B), and it was not until 72 hours that we observed statistically enhanced growth of the *P. aeruginosa* PA14 grown in M9 with 10 and 20 mM N-acetyl-D-glucosamine.

Upon determining that various concentrations of these free glycans had no major effect on bacterial growth *in vitro*, we assessed the effects of free glycans during infection. Free glycans were solubilized in water and added at a concentration of 20 mM to plates to evaluate the role of glycans during infection. The addition of N-acetyl-glucosamine drastically decreased the survival of RNAi control and RNAi *mul-1 C. elegans* survival on *P. aeruginosa* PA14 (Figure 6A). This was expected, based upon previous studies which have shown that N-acetyl-D-glucosamine can alter pathogen virulence characteristics and increase virulence factor expression, specifically pyocyanin (35,36). N-acetyl neuraminic acid (Figure 6B) and D-galactose (Figure 6C) had non-significant effects on RNAi control or RNAi *mul-1* nematode survival. N-acetyl-D-galactosamine supplementation fully restored RNAi *mul-1* nematode survival to RNAi control survival and had no effect on RNAi control nematode survival (Figure 6D), which suggests that *P. aeruginosa* specifically utilizes N-acetyl-D-galactosamine from MUL-1 during infection.

**Figure 6.**
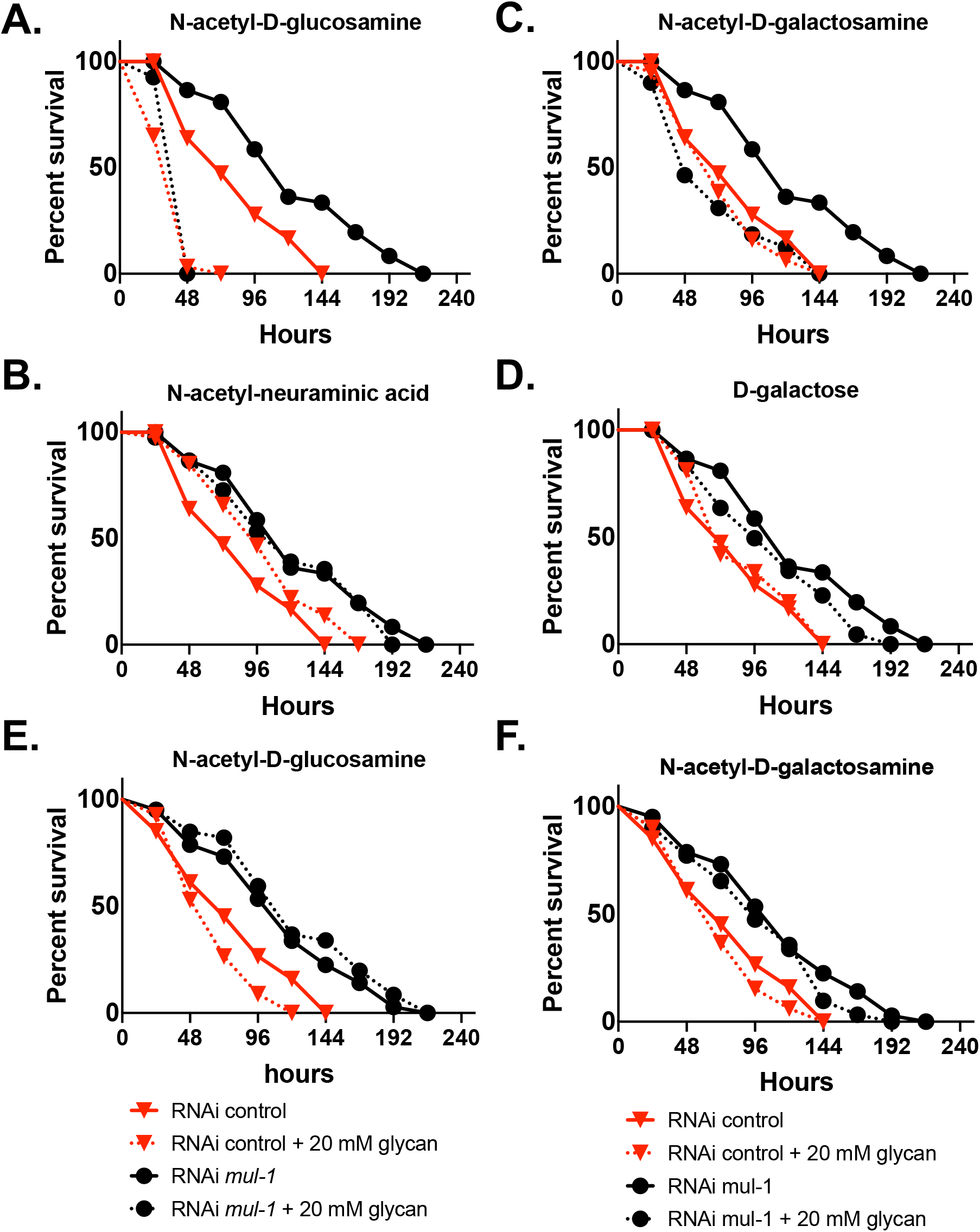
Free glycans in nematode growth medium alters the resistance phenotype of *C. elegans* to *P. aeruginosa* **PA14**. N2 wild-type animals were exposed to two generations of RNAi. Young adult animals were transferred to full lawns of *P. aeruginosa* PA14 on nematode growth media supplemented with **(A)** 20 mM N-acetyl-D-glucosamine, **(B)** 20 mM N-acetyl-neuraminic acid, **(C)** 20 mM N-acetyl-D-galactosamine, or **(D)** 20 mM D-galactose and nematode survival was monitored daily. Animals were considered dead upon failure to respond to touch. Animals missing from the agar plate were censored on the day of loss. Red lines with triangles represent the survival curves of RNAi control animals, without glycans present (solid line) and with glycans present (dotted line). Blue lines with circles represent the survival curves of RNAi *mul-1* animals, without glycans present (solid line) and with glycans present (dotted line). The KaplanMeier method was used to calculate the survival fractions, and statistical significance between survival curves was determined using the log-rank test. 3 biological replicates, 180 total animals per condition. To determine if the glycans needed to be present in the nematode growth medium for *C. elegans* access, *P. aeruginosa* PA14 was grown for 8 hours at 37°C in shaking culture with or without **(E)** 20 mM N-acetyl-D-glucosamine or **(F)** 20 mM N-acetyl-D-galactosamine prior to seeding full lawns of the bacteria. N2 wild-type animals were exposed to two generations of RNAi. Young adult animals were transferred to full lawns of *P. aeruginosa* PA14 on non-supplemented nematode growth medium. Red lines with triangles represent the survival curves of RNAi control animals, without glycans present (solid line) and with glycans present (dotted line). Blue lines with circles represent the survival curves of RNAi *mul-1* animals, without glycans present (solid line) and with glycans present (dotted line). The KaplanMeier method was used to calculate the survival fractions, and statistical significance between survival curves was determined using the log-rank test. 3 biological replicates, 180 total animals per condition.

To determine whether the pathogens must be exposed to the free glycans during infection, or whether the free glycans cause a long-lasting increase in virulence of the pathogens, bacteria were first cultured in the presence of free glycans at concentrations used on the plates used in killing assays. Bacteria were then collected and used to seed full lawn plates, on which survival for the RNAi control and RNAi *mul-1* nematodes was tested. Addition of the free glycans, N-acetyl-D-glucosamine (Figure 6E) and N-acetyl-D-galactosamine (Figure 6F), to the media prior to seeding had no effect on nematode survival on PA14 lawns. This suggests that the free glycans must be present and possibly absorbed by the bacteria as entering the nematode, or absorbed by the nematodes, so that the bacteria have access to the glycans once inside the nematode intestine.

### Free glycans alter binding and internalization of *P. aeruginosa* to human lung epithelial cells

Mucins play conserved roles in maintaining homeostasis and providing protection from infection at epithelial barriers. Human lung epithelial cells express mucins, and some of these have been implicated in the binding and internalization of *P. aeruginosa* into various lung cell lines. Human and mammalian mucins contain many of the same O-linked oligosaccharides; thus, we hypothesized that the free glycans identified in the *C. elegans* system would have similar effects on altering the binding to and cell death of human lung cells.

Free glycans, N-acetyl-D-glucosamine and N-acetyl-D-galactosamine, were tested for their effects on bacterial growth in cell culture media. A range of concentrations of both of the glycans were added to media and bacterial growth was monitored over 24 hours. Within the first 4 hours, the glycans had no significant effect on bacterial growth (Supplemental Figure 4A and 4B). Bacterial binding to lung cells was quantified at time points during which there were no significant changes in bacterial growth. Human lung alveolar epithelial cells, A549 cells, were used to assess bacterial binding in the presence of free glycans because these cells express MUC1 and MUC5a mucins based upon mRNA transcript profiling (37,38). We reasoned that the use of this cell culture model system, with only 2 mucins, would make it simpler to distinguish which specific human mucin is involved in the binding phenotypes observed. The A549 human lung alveolar cells were seeded and grown for approximately 24 hours until 80% confluency. 2 mM free glycans were added to cultures at the same time as bacteria were added. Mid log-phase *P. aeruginosa* PA14 were added to cell cultures at a Multiplicity of Infection (MOI) of 100 and at 4 hours, bacterial binding was assessed. Both N-acetyl-D-glucosamine and N-acetyl-D-galactosamine significantly increased bacterial adherence (Figure 7A).

**Figure 7.**
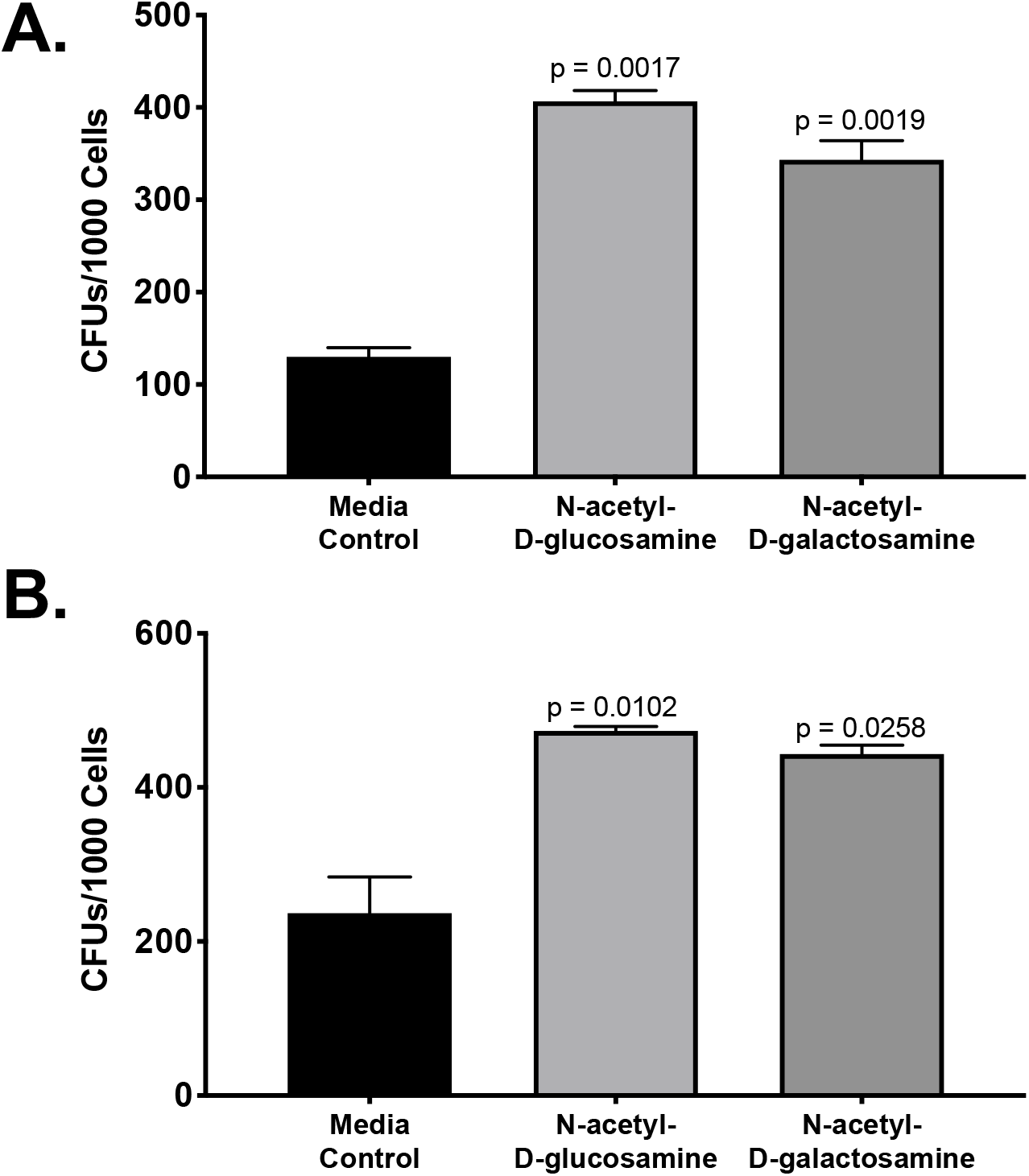
Free glycans alter binding and internalization of *P. aeruginosa* PA14 to A549 human lung cells. **(A)** Approximately 1 × 10^5^ A549 lung cells (~85% confluency) were grown for 2 days with or without 2 mM N-acetyl-D-glucosamine or 2 mM N-acetyl-D-galactosamine supplemented media and infected with an MOI of 100 for 4 hours with *P. aeruginosa* PA14. Intracellular and extracellular bacteria were quantified similarly, as described in methods, but cells were not treated with gentamycin prior to isolating bacteria. Cell viability was measured using *Promega* Cell-Titre Glo 2.0. Serial dilutions were plated to calculate CFUs and normalized to the total number of live cells. 3 biological replicates, each with 3 technical replicates. P values denoted in figure, calculate via the t test. **(B)** Approximately 1 × 10^5^ A549 lung cells (~85% confluency) were grown for 2 days without glycans preset in the media. On the day of the experiment, 2 mM N-acetyl-D-glucosamine or 2 mM N-acetyl-D-galactosamine was added to the media at the same time as the cells were infected with an MOI of 100 for 4 hours with *P. aeruginosa* PA14. Intracellular and extracellular bacteria were quantified similarly, as described in methods, but cells were not treated with gentamycin prior to isolating bacteria. Cell viability was measured using *Promega* Cell-Titre Glo 2.0. Serial dilutions were plated to calculate CFUs and normalized to the total number of live cells. 3 biological replicates, each with 3 technical replicates. P values denoted in figure, calculate via the t test.

To determine if the free glycans are required to be processed by the cells prior to use by *P. aeruginosa*, 2 mM free glycans were added to A549 Lung cells at the time of seeding and remained present for the ~24 hours prior to bacteria were added. *P. aeruginosa* was again added at an MOI of 100 to cells and bacterial binding was measured at 4 hours. When glycans were added at the time of cell seeding, we saw that bacterial binding (Figure 7C) was increased for cells supplemented with both N-acetyl-D-glucosamine and N-acetyl-D-galactosamine.

We have shown that free glycans, specifically N-acetyl-D-galactosamine, can supplement for MUL-1 during *P. aeruginosa* infection of *C. elegans*. In order to determine if these free glycans are able to supplement for mucins in human lung cells, siRNA silencing (lentiviral transfection) was used to knock down expression of human *muc1* in A549 human lung cell model. Knockdown did not result in any growth defects of the cells, but it reduced the number of bacteria bound per cell (Figure 8). This is similar to observations in *C. elegans*, suggesting that mucins and glycans play similar roles in different models of bacterial pathogenesis. Thus, as expected, when N-acetyl-D-glucosamine and N-acetyl-D-galactosamine were added to control and *muc1* siRNA knockdown they enhanced bacterial binding. Both free glycans increased binding of *P. aeruginosa* to the control A549 lung cells when the glycans were added to the media at the same time as the bacteria (Figure 8). The free glycans also reversed the binding defect of *P. aeruginosa* to *muc1* siRNA knockdown cells, restoring the number of bacteria per cell to that of control cells in media alone (Figure 8).

**Figure 8.**
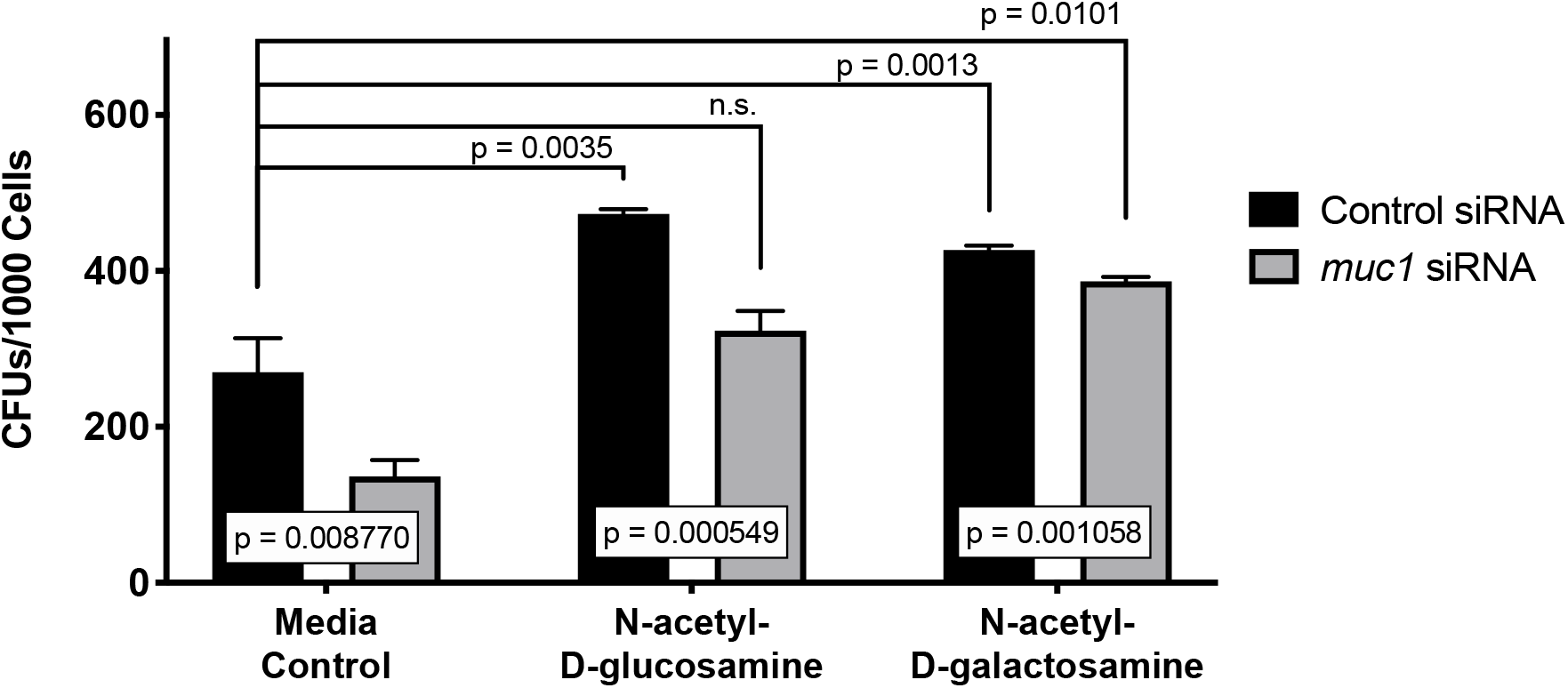
Free glycans reverse the *P. aeruginosa* binding defects to *muc1* siRNA knockdown human lung cells (A549). Sigma MISSION TRC control and *muc1* shRNA Lentiviral Particles were used to create control and *muc1* knockdown cells. Approximately 1 × 10^5^ A549 lung cells (~85% confluency) were grown for 2 days without glycans preset in the media. On the day of the experiment, 2 mM N-acetyl-D-glucosamine or 2 mM N-acetyl-D-galactosamine was added to the media at the same time as the cells were infected with an MOI of 100 for 4 hours with *P. aeruginosa* PA14. Intracellular and extracellular bacteria were quantified, as described in methods. Cells were not treated with gentamycin prior to isolating bacteria. Cell viability was measured using *Promega* Cell-Titre Glo 2.0. Serial dilutions were plated to calculate CFUs and normalized to the total number of live cells. 3 biological replicates, each with 3 technical replicates. P values denoted in figure, calculate via the t test. n.s. is non-significant.

## Discussion

This work details the ability of the pathogen *P. aeruginosa* to use mucin-derived glycans to successfully colonize mucosal epithelial barriers. Our data shows that silencing or deleting a pathogen-induced immune effector gene, *mul-1*, results in enhanced resistance to *P. aeruginosa*. These results were unexpected because we and others have shown that *mul-1* expression is increased upon pathogen infection as part of a general immune response (21,22). Understanding the mechanisms by which *P. aeruginosa* binds to and obtains resources from various mucins may provide a better understanding of host-pathogen interactions and allow for the development of targeted therapeutics.

*P. aeruginosa* is able to utilize mucin-derived glycans to colonize the host, which we have demonstrated by adding free glycans to *C. elegans* growth media during infection. We have shown that one specific glycan, N-acetyl-D-galactosamine, had no effect on control nematodes, but rescued the enhanced resistance phenotype of the *mul-1* RNAi nematodes. This phenomenon appears to be conserved from *C. elegans* to humans, as *P. aeruginosa* binding to human lung epithelial cells was also enhanced in the presence of free glycans, and free glycans reversed the binding defect of *P. aeruginosa* to *muc1* siRNA human lung cells.

In addition to enhanced resistance to *P. aeruginosa* infection, resistance to *S. enterica* is enhanced upon silencing of *mul-1*, which suggests that the use of mucin-derived glycans may be a conserved mechanism by a variety of pathogens that infect mucosal epithelial barriers. In order to fully understand the specific role that *mul-1* plays for pathogens during infection, we tested the ability of *P. aeruginosa* and *S. enterica* to access free glycans from MUL-1 during infection. In order for bacteria to access glycans from mucins, pathogens must first enzymatically cleave the sialic acid cap from the end of polysaccharide chains. Both pathogens have been reported to use free glycans as carbon sources, but only *S. enterica* encodes and expresses a bacterial sialidase (28,30). *C. elegans* encodes an enzyme, *ctsa-1.1*, that is predicted to perform a similar enzymatic function as bacterial sialidases, has a predicted secretion signal, and is expressed in the intestine. We have shown that RNAi silencing of *ctsa-1.1* results in enhanced resistance to *P. aeruginosa* but not enhanced resistance to *S. enterica*. This suggests that *S. enterica* is able to use its bacterial-encoded sialidase to gain access to glycans, but *P. aeruginosa* cannot access glycans without the *C. elegans* enzyme. It is possible that other intestinal mucins serve similar roles in binding and acting as a carbon source for pathogens, which may depend upon the composition of the glycan chains of the mucin and the specific use of the glycans by the pathogens.

One of the most interesting implications of our data is that while *mul-1* expression is enhanced upon *C. elegans* infection with *P. aeruginosa* and *S. enterica*, the animals would apparently be at an advantage without *mul-1* during infection with *P. aeruginosa* and *S. enterica*. The bacteria are able to access the glycans present on the glycan chains by using either host-expressed or bacterial expressed sialidase enzymes. These glycans aid in the colonization of the host, suggesting that despite the fact that MUL-1 in an innate immune response factor, pathogens have evolved to use this mucin and the glycans that decorate the peptide. The ability to control a pathogen’s access to mucins and the provided glycans could prove to be a method to prevent and/or disrupt infection. In certain disease states, opportunistic pathogens are able to colonize and establish an infection when there are changes in the levels of mucus and expression levels of mucins. In some cases, only the glycosylation patterns of mucins change and this provides enhanced binding for pathogens. If there were therapeutics available to prevent binding of pathogens to mucin oligosaccharides or to control mucin expression, this would be extremely beneficial in treating disease and could prevent opportunistic pathogen infection.

## Acknowledgements

None.

## Methods

### Bacterial Strains

The following bacterial strains were used: *Escherichia coli* OP50, *E. coli* HT115 (DE3), *Pseudomonas aeruginosa* PA14, *P. aeruginosa* PA14-GFP, *Salmonella enterica* serovar Typhimurium 1344. All strains were grown on Luria-Bertani (LB) agarose plates or LB broth at 37°C shaking at 250 RPM.

### C. elegans Strains and Growth Conditions

*C. elegans* hermaphrodites were maintained on *E. coli* OP50 at 20°C unless otherwise indicated. Bristol N2 was used as the wild-type control strain. *mul-1* (*ac7*) CRISPR/Cas9 ~1650 bp deletion mutant of isoform A and B (565bp and 952bp deleted, with generated termination codon); predicted truncated protein has 46 amino acids. Gut-sensitive RNAi lines: MGH171 alxIs9 [vha-6p∷sid-1∷SL2∷GFP]; kbIs7 [nhx-2p∷rde-1 + rol-6(su1006)]. -- were obtained from the Caenorhabditis Genetics Center (University of Minnesota, Minneapolis, MN).

### RNA Interference (RNAi)

RNAi was used to generate loss-of-function RNAi phenotypes by feeding nematodes *E. coli* strain HT115 (DE3) expressing double stranded RNA (dsRNA) homologous to a target gene (39–41). RNAi was carried out as described previously (42). Briefly, *E. coli* with appropriate vectors were grown in LB broth containing ampicillin (100 μg/mL) and tetracycline (12.5 μg/mL) at 37°C overnight and plated onto NGM plates containing 100 mg/mL ampicillin and 6 mM isopropyl b-D-thiogalactoside (IPTG) (RNAi plates). RNAi-expressing bacteria were allowed to grow overnight at 37°C. Gravid adults were transferred to RNAi-expressing bacterial lawns and allowed to lay eggs for 8 hours. The gravid adults were removed, and the eggs were allowed to develop at 20°C to young adults for subsequent assays. *unc-22* RNAi was included as a positive control to account for the RNAi efficiency. All RNAi clones except *mul-1* were from the Ahringer RNAi library (Open BioSource). The *mul-1* clone was obtained from the Vidal RNAi library (Open BioSource).

### Killing Assay, Survival on Pathogens

Bacterial lawns were prepared by inoculating individual bacterial colonies into 2 mL of LB with 50 mg/mL kanamycin and growing them for 7-8 hours on a shaker at 37°C. For the colonization assays, bacterial lawns of *P. aeruginosa* or *S. enterica* ser Typhimurium were prepared by spreading 35 μL of the culture over the complete surface of 3.5-cm-diameter modified NGM agar plates (3.5% instead of 2.5% peptone). Young adult animals were transferred to full lawns of *P. aeruginosa* PA14 or *S. enterica* ser Typhimurium and nematode survival was monitored daily. Animals were considered dead upon failure to respond to touch. Animals missing from the agar plate were censored on the day of loss. The KaplanMeier method was used to calculate the survival fractions, and statistical significance between survival curves was determined using the log-rank test.

### Avoidance Assay on Pathogens

The bacterial lawns were prepared by inoculating individual bacterial colonies into 2 mL of the corresponding broth mentioned above and growing them for 7-8 hours on a shaker at 37°C. 20 μL of the culture was plated onto the center of a 3.5-cm plate and incubated at 37°C for 12-16 hours. For *P. aeruginosa* PA14, modified NGM (3.5% instead of 2.5% peptone) plates were used. Thirty synchronized young gravid adult hermaphroditic animals grown on *E. coli* HT115 (DE3) containing control vector or an RNAi clone targeting a gene were transferred outside the bacterial lawns. The numbers of animals on and off the lawns were counted at the indicated times for each experiment. Three 3.5-cm plates were used per trial in each experiment. Experiments were performed at 25°C. The percent occupancy was calculated as (# on lawn/# total). At least three independent experiments were performed.

### *P. aeruginosa*-GFP Colonization Assay

Bacterial lawns were prepared by inoculating individual bacterial colonies into 2 mL of LB with 50 mg/mL kanamycin and growing them for 7-8 hours on a shaker at 37°C. For the colonization assays, bacterial lawns of *P. aeruginosa*-GFP were prepared by spreading 35 μL of the culture over the complete surface of 3.5-cm-diameter modified NGM agar plates (3.5% instead of 2.5% peptone). The plates were incubated at 37°C for 12-16 hours and then cooled to room temperature for at least 1 h before seeding with young gravid adult hermaphroditic animals. The assays were performed at 25°C. At the indicated times for each experiment, the animals were transferred from *P. aeruginosa*-GFP plates to fresh E. coli OP50 plates and visualized within 5 minutes under a fluorescence microscope.

### Quantification of Intestinal Bacterial Loads

*P. aeruginosa*-GFP lawns were prepared as described above. For quantification of colony forming units (CFU) at various timepoints, bacterial lawns of *P. aeruginosa*-GFP were prepared by spreading 35 μL of the culture over the complete surface of 3.5-cm-diameter modified NGM agar plates (3.5% instead of 2.5% peptone). The plates were incubated at 37°C for 12-16 hours and then cooled to room temperature for at least 1 hour before seeding with young adult hermaphroditic animals. The assays were performed at 25°C. At indicated times for each experiment, the animals were transferred *from P. aeruginosa*-GFP plates to the center of fresh *E. coli* plates for 30 min to eliminate *P. aeruginosa*-GFP stuck to their body. Animals were transferred again to the center of a new *E. coli* plate for 30 additional minutes to further eliminate external *P. aeruginosa*-GFP. Afterward, ten animals/condition were transferred into 50 μL of PBS plus 0.01% Triton X-100 and ground using glass beads. Serial dilutions of the lysates (10^−1^, 10^−2^, 10^−3^, 10^−4^, 10^−5^) were seeded onto LB plates containing 50 mg/mL of kanamycin to select for *P. aeruginosa*-GFP cells. Plates were incubated overnight at 37°C. Single colonies were counted the following day and are represented as the number of bacterial cells or CFUs per animal. Three independent experiments were performed for each condition.

### Fluorescence Imaging

Fluorescence imaging was carried out as described previously (42). Briefly, animals were anesthetized using an M9 salt solution containing 30 mM sodium azide and mounted onto 2% agar pads. The animals were then visualized using a Leica M165 FC fluorescence stereomicroscope.

### RNA Isolation and Quantitative Reverse Transcription-PCR (qRT-PCR)

Animals were synchronized by egg laying. Approximately 35 N2 gravid adult animals were transferred to 10-cm RNAi plates seeded with *E. coli* HT115 (DE3) expressing the appropriate vectors and allowed to lay eggs for 8 hours. Gravid adults were then removed, and the eggs were allowed to develop at 20°C. For gene expression analysis in the *P. aeruginosa* infection assays, the animals were first grown on 10-cm RNAi plates seeded with *E. coli* HT115 expressing either the empty vector RNAi control or *mul-1* RNAi until the young adult stage. Subsequently, the animals were collected, washed with M9 buffer, and transferred to 10-cm modified NGM plates (3.5% instead of 2.5% peptone) seeded with 300 μL of *P. aeruginosa* culture grown overnight. The *P. aeruginosa* plates were prepared by spreading 300 μL of the *P. aeruginosa* culture on the surface of the modified NGM plates, followed by an overnight incubation at 37°C. After transfer of the animals, the *P. aeruginosa* plates were incubated at 25°C for 8 hours. Animals on the control *E. coli* plates were also incubated at 25°C. After the desired treatment, the animals were collected, washed with M9 buffer, and frozen in TRIzol reagent (Life Technologies, Carlsbad, CA). Total RNA was extracted using the RNeasy Plus Universal Kit (Qiagen, Netherlands). Residual genomic DNA was removed using TURBO DNase (Life Technologies, Carlsbad, CA). A total of 6 μg of total RNA was reverse-transcribed with random primers using the High-Capacity cDNA Reverse Transcription Kit (Applied Biosystems, Foster City, CA). qRT-PCR was conducted using the Applied Biosystems One-Step Real-time PCR protocol using SYBR Green fluorescence (Applied Biosystems) on an Applied Biosystems 7900HT real-time PCR machine in 96-well-plate format. 25 μL reactions were analyzed as outlined by the manufacturer (Applied Biosystems). The relative fold-changes of the transcripts were calculated using the comparative CT^(2−ΔΔCT)^ method and normalized to pan-actin (*act-1*, -*3*, -*4*). The cycle thresholds of the amplification were determined using StepOnePlus software (Applied Biosystems). All samples were run in triplicate. The primer sequences are available in Table S1.

### *C. elegans* Longevity Assays, Cultivation of *C. elegans* on Heat-Killed*E. coli* OP50

A single colony of *E. coli* OP50 was inoculated in 100 mL of LB broth in a 500 mL Erlenmeyer flask and incubated at 37°C at 225 rpm shaking for 24 hours. Bacteria were concentrated 20 times and heat-killed at 100°C for 1 hour. Bacterial death was confirmed by failure to grow on LB plates at 37°C overnight. The concentrated, heat-killed bacteria were seeded on NGM plates containing 50 mg/mL of kanamycin and 100 mg/mL of streptomycin. Young adult wild-type N2 animals grown on *E. coli* HT115 RNAi control or target gene RNAi plates were washed with M9 medium and incubated at room temperature for 1 hour with M9 medium containing 50 mg/mL of kanamycin to remove live bacteria from their intestinal lumen. The animals were then washed with M9 medium and transferred to NGM plates containing heat-killed *E. coli* OP50 and incubated at 20°C for the duration of the assay. Remaining animals were transferred when the heat-killed *E. coli* OP50 lawn was reduced.

### *C. elegans* Longevity Assays, Cultivation of *C. elegans* on Heat-Killed *E. coli* HT115 (DE3) RNAi

Lifespan assays were performed on RNAi plates containing *E. coli* HT115 (DE3) with the appropriate vector. Animals were synchronized on RNAi plates and incubated at 20°C. At the L4 larval stage, the animals were transferred onto the corresponding RNAi plates. Animals were scored on a daily basis as alive, dead, or gone. Animals that failed to display touch-provoked movement were scored as dead. Experimental groups contained 60 to 100 animals. Young adult animals were considered as day 0 for the lifespan analysis. The assays were performed at 20°C. Three independent experiments were performed.

### Bacterial Growth Assays

Individual bacterial colonies were inoculated into 2 mL of LB and grown overnight, shaking at 225 RPM at 37°C. Overnight cultures were diluted to an OD_570_ of 0.05 in either LB media, M9 Media, or DMEM F12+K + 10% Heat-Inactivated FBS. 100 μL bacterial cultures were placed in individual wells of a 96 well plate with various glycans added at indicated concentrations. Bacterial growth was monitored over time by measuring the OD_570_ at indicated time points.

### Cell Lines

The A549 human type II alveolar epithelial cell line (ATCC # CCL-185, provided by David Lewinsohn’s lab at Oregon Health and Science University); passage numbers 3-10 were used. A549 cells were maintained in Ham’s F-12K (Kaighn’s) Medium (Gibco 21127022) supplemented with 10% heat-inactivated fetal bovine serum (SAFC Biosciences, Lenexa, KS) without antibiotics. Cells were grown at 37°C with 5% CO_2_ and seeded every 4 days when confluency was approximately 85%.

### Lentiviral Transduction with MISSION TRC shRNA Lentivirus Particles

The MISSION TRC shRNA Lentiviral Transfection protocol was used. 1 × 10^6^ A549 cells were plated in 60 mm dishes and grown 20 hours at 37°C 5% CO_2_ to ~60-70% confluency. Media was replaced without 8 μg/mL Hexadimethrine bromide. An MOI of ~0.5 (~13 μL of stock lentivirus) was added to plates and incubated for 6 hours. Media was replaced without antibiotics for 24 hours. The following day, media was replaced with medium containing 5 μg/mL puromycin. Media was replaced with fresh puromycin containing media every 3-4 days until resistant colonies grew to confluency. Cells were then used for indicated assays.

### Gentamycin Protection Assay – Bacterial binding to cells

The gentamycin protection assay protocol was used, as previously described (43), with some modifications (44). A549 cells were grown in 96-well plates with or without free glycans present for 2 days at 37°C, 5% CO_2_ until they reached approximately 85% confluency, corresponding to 1 × 10^5^ cells per well. Wells were washed four times with sterile PBS and serum-free DMEM F12 was added for 2 h. *P. aeruginosa* PA14 was grown overnight in LB at 37°C shaking, diluted 1:10 and allowed to grow under the same conditions for 4 hours (obtaining mid-log phase bacteria). A549 cells were infected with *P. aeruginosa* PA14 at an MOI of 100 and incubated for 2 hours at 37°C, 5% CO_2_. From the original bacterial suspension, serial dilutions were plated were prepared to verify the starting concentration of *P. aeruginosa*. Following incubation, the supernatants were removed, the wells were washed four times with PBS, and the numbers of associated or internalized bacteria were assessed as follows: 30 μL of 0.05% trypsin-EDTA was added to each well, incubated for 2 min at 37°C. Cells were lysed with 70 μL of 0.1% Triton X-100 for 2 min at 37°C. Lysates were removed and serial dilutions were plated. Cell viability was measured using *Promega* Cell-Titre Glo 2.0 reagent (quantitates cellular ATP levels) and CFUs were normalized per cell under each condition. These results represent the number of both Attached and Intracellular bacteria (CFU / # of cells). To quantify internalized bacteria, after the 4 hour incubation with *P. aeruginosa* PA14 and four PBS washes, 200 μg/mL gentamicin sulfate in 50 μL of medium was added to each well and incubated for 90 minutes to eliminate extracellular bacteria. After the incubation, supernatants were removed, and serial dilutions were plated to quantitate the number of extracellular *P. aeruginosa* PA14 that may have survived gentamycin treatment. A549 cells were then washed four times with PBS, trypsinized, and lysed as described above; serial dilutions were performed and plated onto LB agar plates. Again, CFUs were normalized to viable cells. The results represent the number of intracellular bacteria (CFU / # of cells).

### Quantification and Statistical Analysis

Statistical analysis was performed with Prism 7 (GraphPad). All error bars represent the standard deviation (SD). The two-sample t test was used when needed, and the data were judged to be statistically significant when p < 0.05. In the figures, asterisks (*) denotes statistical significance as follows: *, p < 0.05, **, p < 0.001, ***, p < 0.0001, as compared to the appropriate controls. The KaplanMeier method was used to calculate the survival fractions, and statistical significance between survival curves was determined using the log-rank test. All experiments were performed in triplicate.

---

**Figure S1. RNAi silencing of several intestine-expressed mucins alters the resistance phenotype to *P. aeruginosa* PA14. (A)** N2 wild-type animals were exposed to two generations of RNAi. Young adult animals were transferred to full lawns of *P. aeruginosa* PA14 and nematode survival was monitored daily. Animals were considered dead upon failure to respond to touch. Animals missing from the agar plate were censored on the day of loss. The KaplanMeier method was used to calculate the survival fractions, and statistical significance between survival curves was determined using the log-rank test. 3 biological replicates, 180 total animals per condition. **(B)** Because RNAi for *let-653* is larval lethal, L4 larval stage N2 wild-type animals were transferred to full lawns of both control and *let-653 E. coli* HT115(DE3) RNAi bacteria. Young adults were transferred to full lawns of *P. aeruginosa* PA14 and nematode survival was monitored daily. Animals were considered dead upon failure to respond to touch. Animals missing from the agar plate were censored on the day of loss. The KaplanMeier method was used to calculate the survival fractions, and statistical significance between survival curves was determined using the log-rank test. 3 biological replicates, 120 total animals per condition. **(C)** Time (hours) to 50% death upon *P. aeruginosa* PA14 exposure was calculated for each of the survival curves in (A) and (B) using *Graphpad Prism 8 software* and is reported as TD_50_.

**Figure S2. The ability of *S. enterica* ser Typhimurium to access O-linked glycans provided by *mul-1* in the intestine alters the resistance phenotype of *C. elegans* nematodes. (A)** N2 wild-type animals were exposed to two generations of RNAi targeting *mul-1*, and *F41C3.5*. Young adult animals were transferred to full lawns of *S. enterica* ST1334 and nematode survival was monitored daily. Animals were considered dead upon failure to respond to touch. Animals missing from the agar plate were censored on the day of loss. The KaplanMeier method was used to calculate the survival fractions, and statistical significance between survival curves was determined using the log-rank test. 3 biological replicates, 180 total animals per condition. **(B)** After RNAi, young adult nematodes were transferred to full lawns of *S. enterica* ST1334-GFP (kan^r^). At 24 hours post *S. enterica* ST1334 exposure, nematodes were transferred to fresh *E. coli* OP50 lawns to remove *S. enterica*. Worms were ground and serial dilutions were plated on LB-kanamycin plates to calculate Colony Forming Units (CFUs) per worm (nematode). 3 biological replicates, 90 total animals per condition. **(C)** After RNAi, young adult nematodes were transferred to full lawns of *P. aeruginosa* PA14-GFP (kan^r^). At 24 hours post *S. enterica* ST1334 exposure, nematodes were transferred to fresh *E. coli* OP50 lawns to remove *S. enterica*. Worms were then transferred to fresh *E. coli* OP50 plates and at indicated time points, nematodes were ground and serial dilutions were plated on LB-kanamycin plates to calculate persistant Colony Forming Units (CFUs) per worm (nematode). 3 biological replicates, 90 total animals per condition

**Figure S3. Free glycans have limited effects on *P. aeruginosa* growth in bacterial growth media.** Individual bacterial colonies were inoculated into 2 mL of LB and grown overnight, shaking at 225 RPM at 37°C. Overnight cultures were diluted to an OD_600_ of 0.05 in either **(A)** LB media or **(B)** M9 Media supplemented with varying concentrations of free glycans. 100 μL bacterial cultures were placed in individual wells of a 96 well plate with various glycans added at indicated concentrations. Bacterial growth was monitored over time by measuring the OD_600_ at indicated time points.

**Figure S4. Free glycans have limited effects on *P. aeruginosa* growth in cell growth media.** Individual bacterial colonies were inoculated into 2 mL of LB and grown overnight, shaking at 225 RPM at 37°C. Overnight cultures were diluted to an OD_600_ of 0.05 in DMEM F12+K + 10% Heat-Inactivated FBS with varying concentrations of **(A)** N-acetyl-D-glucoasmine or **(B)** N-acetyl-D-galactosamine. 100 μL bacterial cultures were placed in individual wells of a 96 well plate with various glycans added at indicated concentrations. Bacterial growth was monitored over time by measuring the OD_600_ at indicated time points.

