## Supplementary Files for "Host mucin is subverted by *Pseudomonas aeruginosa* during infection to provide free glycans required for successful colonization"

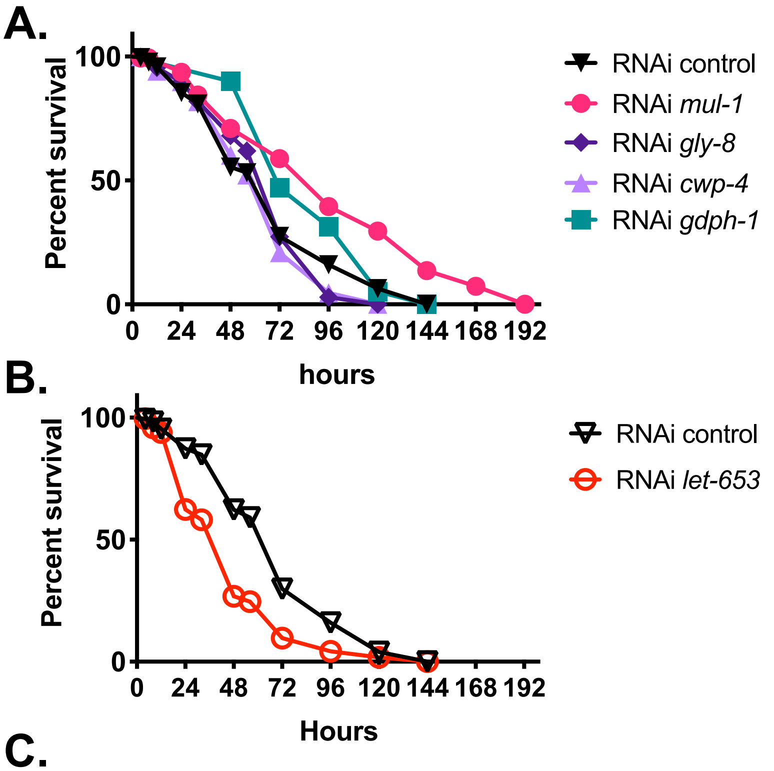


C.

| RNAi Condition | TD50 (hours) |
|---|---|
| RNAi Control | 54.6 |
| RNAi *mul-1* | 82.8 |
| RNAi *gdph-1* | 73.2 |
| RNAi *gly-8* | 59.4 |
| RNAi *cwp-4* | 56 |
| RNAi *let-653* | 38.6 |

**Figure S1. RNAi silencing of several intestine-expressed mucins alters the resistance phenotype to *P. aeruginosa* PA14. (A)** N2 wild-type animals were exposed to two generations of RNAi. Young adult animals were transferred to full lawns of *P. aeruginosa* PA14 and nematode survival was monitored daily. Animals were considered dead upon failure to respond to touch. Animals missing from the agar plate were censored on the day of loss. The KaplanMeier method was used to calculate the survival fractions, and statistical significance between survival curves was determined using the log-rank test. 3 biological replicates, 180 total animals per condition. **(B)** Because RNAi for *let-653* is larval lethal, L4 larval stage N2 wild-type animals were transferred to full lawns of both control and *let-653 E. coli* HT115(DE3) RNAi bacteria. Young adults were transferred to full lawns of *P. aeruginosa* PA14 and nematode survival was monitored daily. Animals were considered dead upon failure to respond to touch. Animals missing from the agar plate were censored on the day of loss. The KaplanMeier method was used to calculate the survival fractions, and statistical significance between survival curves was determined using the log-rank test. 3 biological replicates, 120 total animals per condition. **(C)** Time (hours) to 50% death upon *P. aeruginosa* PA14 exposure was calculated for each of the survival curves in (A) and (B) using *Graphpad Prism 8 software* and is reported as $TD_{50}$.

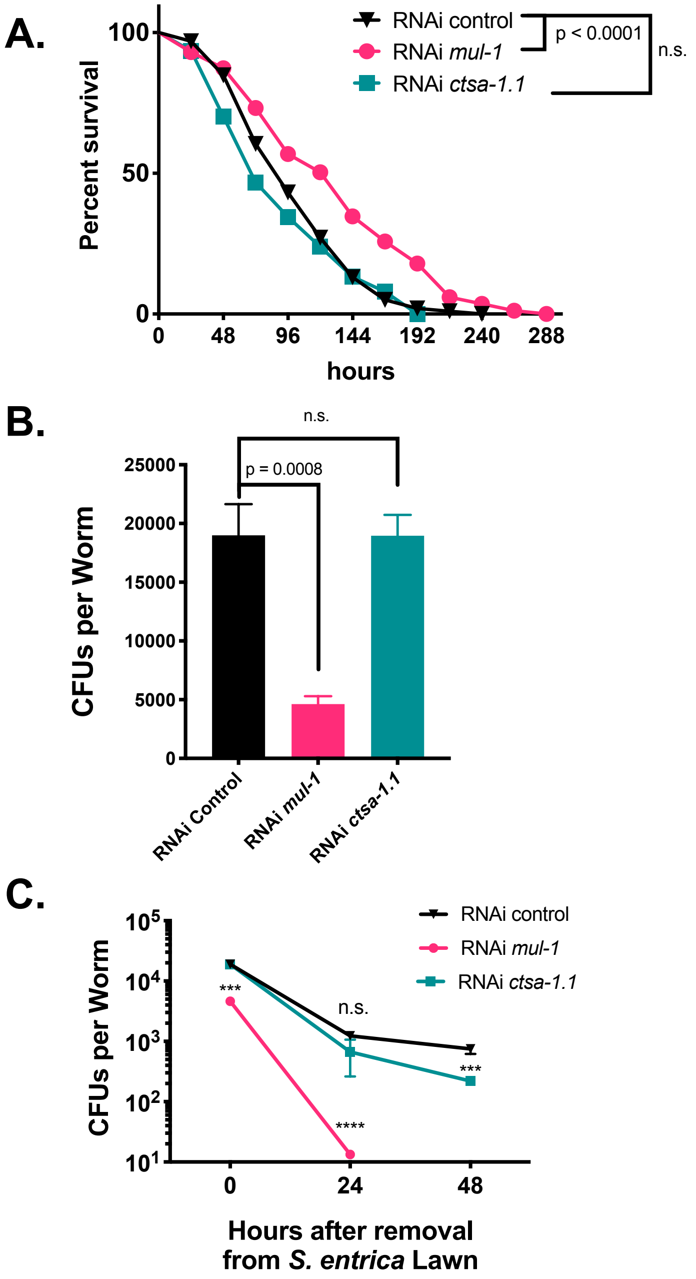


**Figure S2. The ability of *S. enterica* ser Typhimurium to access O-linked glycans provided by *mul-1* in the intestine alters the resistance phenotype of *C. elegans* nematodes. (A)** N2 wild-type animals were exposed to two generations of RNAi targeting *mul-1*, and *F41C3.5.* Young adult animals were transferred to full lawns of *S. enterica* ST1334 and nematode survival was monitored daily. Animals were considered dead upon failure to respond to touch. Animals missing from the agar plate were censored on the day of loss. The KaplanMeier method was used to calculate the survival fractions, and statistical significance between survival curves was determined using the log-rank test. 3 biological replicates, 180 total animals per condition. **(B)** After RNAi, young adult nematodes were transferred to full lawns of *S. enterica* ST1334-GFP (kan$^r$). At 24 hours post *S. enterica* ST1334 exposure, nematodes were transferred to fresh *E. coli* OP50 lawns to remove *S. enterica.* Worms were ground and serial dilutions were plated on LB-kanamycin plates to calculate Colony Forming Units (CFUs) per worm (nematode). 3 biological replicates, 90 total animals per condition. **(C)** After RNAi, young adult nematodes were transferred to full lawns of *P. aeruginosa* PA14-GFP (kan$^r$). At 24 hours post *S. enterica* ST1334 exposure, nematodes were transferred to fresh *E. coli* OP50 lawns to remove *S. enterica.* Worms were then transferred to fresh *E. coli* OP50 plates and at indicated time points, nematodes were ground and serial dilutions were plated on

LB-kanamycin plates to calculate persistant Colony Forming Units (CFUs) per worm (nematode). 3 biological replicates, 90 total animals per condition

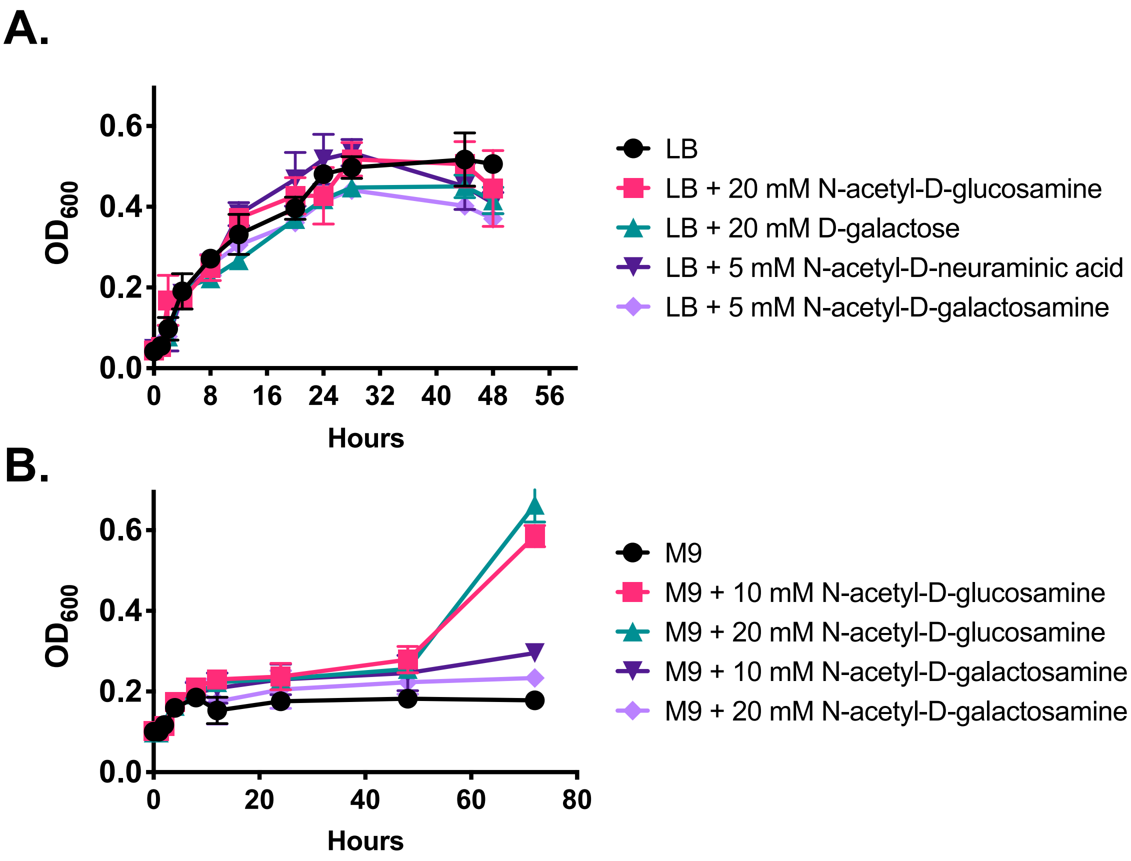


**Figure S3. Free glycans have limited effects on *P. aeruginosa* growth in bacterial growth media.** Individual bacterial colonies were inoculated into 2 mL of LB and grown overnight, shaking at 225 RPM at 37ºC. Overnight cultures were diluted to an $OD_{600}$ of 0.05 in either **(A)** LB media or **(B)** M9 Media supplemented with varying concentrations of free glycans. 100 µL bacterial cultures were placed in individual wells of a 96 well plate with various glycans added at indicated concentrations. Bacterial growth was monitored over time by measuring the $OD_{600}$ at indicated time points.

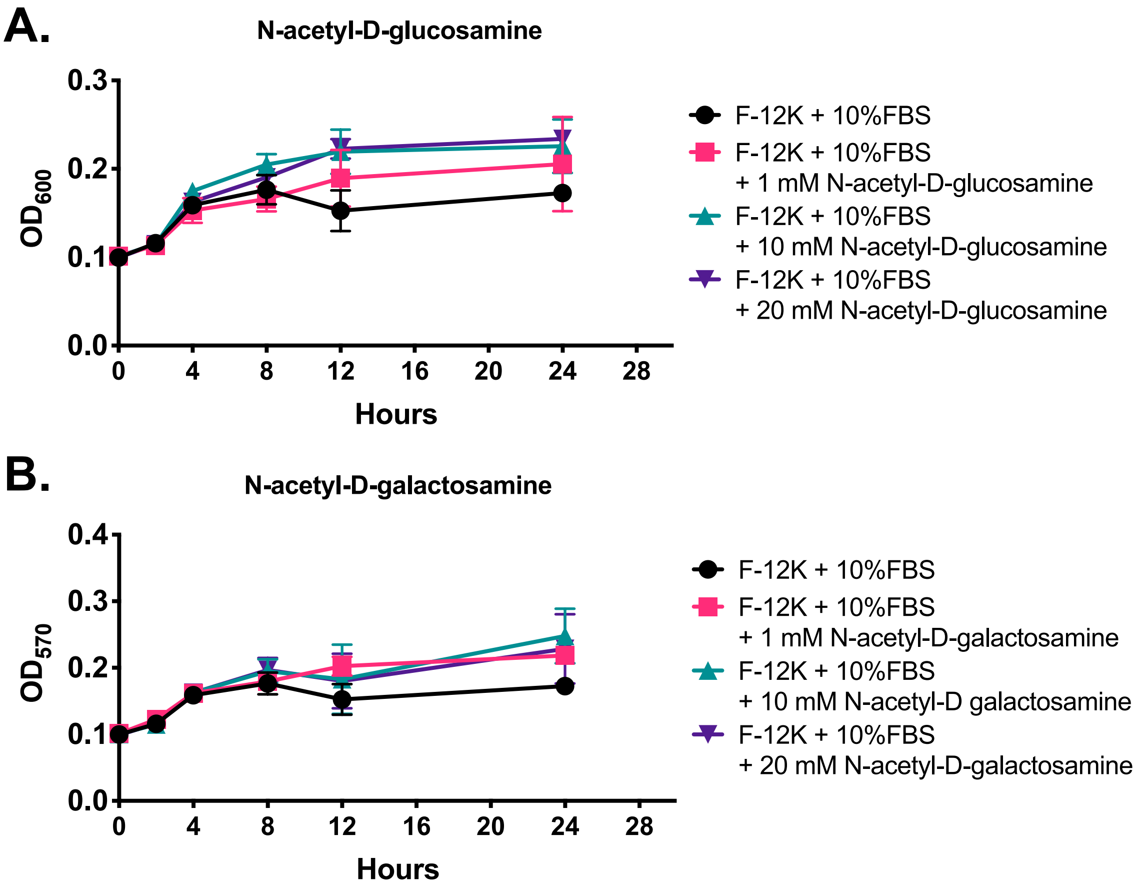


**Figure S4. Free glycans have limited effects on *P. aeruginosa* growth in cell growth media.** Individual bacterial colonies were inoculated into 2 mL of LB and grown overnight, shaking at 225 RPM at 37ºC. Overnight cultures were diluted to an $OD_{600}$ of 0.05 in DMEM F12+K + 10% Heat-Inactivated FBS with varying concentrations of **(A)** N-acetyl-D-glucoasmine or **(B)** N-acetyl-D-galactosamine. 100 µL bacterial cultures were placed in individual wells of a 96 well plate with various glycans added at indicated concentrations. Bacterial growth was monitored over time by measuring the $OD_{600}$ at indicated time points.

**Table S1. Intestine-expressed mucin and mucin enzyme genes found in the *C. elegans* genome.** Because mucins exhibit little sequence similarity at the DNA and amino acid level. *C. elegans* mucins were identified based on conserved serine- and threonine-rich regions subject to heavy O-glycosylation and cysteine-rich regions subject to heavy N-glycosylation. Predicted mucins were further refined based upon mRNA expression in the intestine and changes in gene expression associated with pathogen infection. Selected mucins and proteins associated with glycosylation of mucins are highlighted in Table 1.

| Gene | Sequence | Gene Details | Associated Phenotypes |
|---|---|---|---|
| *let-653* | *C29E6.1* | mucin-like protein similar to highly glycosylated mucins of the apical surface of epithelia | Larval lethal |
| *gpdh-1* | *F47G4.3* | encodes enzyme glycerol 3-phosphate dehydrogenase | Human ortholog associated with Brugada Syndrome 2 |
| *gly-8* | *Y66A7A.6* | encodes predicted transmembrane polypeptide N-acetylgalactosaminyl transferase (ppGaNTase) | none |
| *mul-1* | *F49F1.6* | encodes mucin-like protein containing signal sequence and several ShK toxin domains | Increased upon radiation exposure |
| *cwp-4* | *K11D12.1* | encodes a protein with similarity to mucins, predicted to be secreted | Higher expression in males. Increased upon NaCl exposure |

**Table S2.** Primers used for quantitative Real Time-PCR

| Gene name | Forward primer sequence (5'-3') | Reverse primer sequence (5’-3’) |
|---|---|---|
| Pan-*act* | TCGGTATGGGACAGAAGGAC | CATCCCAGTTGGTGACGATA |
| *mul-1* | GTGATCAGGGATTTGTGCAGAGTC | GATACTTCACATCCGTCTTGCAC |
| *let-653* | CTGTCTCGTGAGAATATGTCC | TTCCACGTCGTCGCATGT |
| *gly-8* | ATGGTTGGAGCCACTGCTAC | TGAACGTGAATCCCCAGTCG |
| *gpdh-1* | AGGCGACAATCGGGTGTAAG | GCACCATGTTCCCCAACAAC |
| *cwp-4* | AAGCGAACAACACCAATGCC | GTCGTTGTCGTCGTAGTGGT |
